# Novel amiloride derivatives that inhibit bacterial motility across multiple strains and stator types

**DOI:** 10.1101/2021.04.13.439105

**Authors:** MI Islam, JH Bae, T Ishida, P Ridone, J Lin, MJ Kelso, Y Sowa, BJ Buckley, MAB Baker

## Abstract

The bacterial flagellar motor (BFM) is a protein complex that confers motility to cells and contributes to survival and virulence. The BFM consists of stators that are ion-selective membrane protein complexes and a rotor that directly connects to a large filament, acting as a propeller. The stator complexes couple ion transit across the membrane to torque that drives rotation of the motor. The most common ion gradients that drive BFM rotation are protons (H^+^) and sodium ions (Na^+^). The sodium-powered stators, like those in the PomAPomB stator complex of Vibrio spp, can be inhibited by sodium channel inhibitors, in particular, by phenamil, a potent and widely used inhibitor. However, relatively few new sodium-motility inhibitors have been described since the discovery of phenamil. In this study, we characterised two possible motility inhibitors HM2-16F and BB2-50F from a small library of previously reported amiloride derivatives. We used three approaches: effect on rotation of tethered cells, effect on free swimming bacteria and effect on rotation of marker beads. We showed that both HM2-16F and BB2-50F stopped rotation of tethered cells driven by Na^+^ motors comparable to phenamil at matching concentrations, and could also stop rotation of tethered cells driven by H^+^ motors. Bead measurements in presence and absence of stators confirmed that the compounds did not inhibit rotation via direct association with the stator, in contrast to the established mode of action of phenamil. Overall, HM2-16F and BB2-50F stopped swimming in both Na^+^ and H^+^ stator types, and in pathogenic and non-pathogenic strains.

**Importance:** Here we characterised two novel amiloride derivatives in the search for antimicrobial compounds that target bacterial motility. Our two compounds were shown to inhibit flagellar motility at 10 μM across multiple strains, from non-pathogenic *E. coli* with flagellar rotation driven by proton or chimeric sodium-powered stators, to proton-powered pathogenic *E. coli* (EHEC/UPEC) and lastly in sodium-powered *Vibrio alginolyticus*. Broad anti-motility compounds such as these are important tools in our efforts control virulence of pathogens in health and agricultural settings.

## Introduction

Motility is essential for many bacteria to find nutrients and escape unfavourable conditions (1). Bacterial cells move through liquids or over moist surfaces using rotating flagella, which are driven by the bacterial flagellar motor (BFM) (2). In peritrichous bacteria such as *E. coli*, when the majority of BFMs rotate counter-clockwise, bacteria move linearly (i.e., run) and when one or more BFM rotates clockwise, the helical filament-bundle unravels and bacteria change direction (i.e., tumble) (3). The BFM contains the stator proteins (e.g., MotA and MotB in *E. coli* and PomA and PomB in *Vibrio* spp.), which form an ion transporter on the inner cell membrane, and the rotor proteins (e.g., FliG, FliM, and FliN), that are driven to rotate by torque generation between the stator and rotor (4). Rotation of the BFM is powered by translocation of specific ions (e.g., H^+^ in *E. coli* and Na^+^ in *Vibrio* spp.) across the cell membrane through the stator complex (4).

Beside locomotion, motility is crucial during the initial infection and colonisation processes of different pathogenic bacteria (5). Locomotion reduces repulsion between the bacterial cell wall and host tissues, promoting attachment to host cells (6). The importance of motility as a virulence factor was first noted in *Campylobacter jejuni*, when only the motile strain of *C. jejuni* was recovered from an infection site where a mixture of motile and non-motile *C. jejuni* phase variants were used to initiate infection (7). It has since been demonstrated that flagellum-mediated motility promotes the virulence of several gram-negative pathogens, such as for *Bordetella bronchiseptica, B. pertussis* (8), *Helicobacter sp*. (9), *Legionella pneumophila* (10), pathogenic *E. coli* strains (11), *Pseudomonas aeruginosa* (12), *Salmonella typhimurium* (13), and *Vibrio cholerae* (14). Aside from host attachment, flagella also contribute to the formation of biofilms by multiple bacterial pathogens, which promote the development of persistent and chronic infections and antibiotic resistance (1).

Amiloride is a diuretic drug and well-characterised inhibitor of the Na^+^ bacterial flagellar motor (15). Amiloride inhibits rotation of the sodium-powered flagellar motor by blocking the translocation of Na^+^ ions through the channel in a competitive and rapidly reversible manner (16). However, amiloride is of limited utility due to its weak anti-motility activity and adverse effects on other metabolic processes (17). The *N*-phenyl substituted amiloride analogue, phenamil, has been identified as a better inhibitor of sodium-powered flagellar motors (17). Phenamil is markedly more potent than amiloride and shows non-competitive inhibition of bacterial motility with slow dissociation (18). Phenamil stops flagellar rotation by binding to a site predicted to lie at the cytoplasmic face of the PomAPomB stator complex (19). Phenamil is a useful tool for flagellar research, but requires high concentrations (>20 μM) and resistant stators can arise from a single point mutation (20, 21). More potent and specific compounds may assist research into stator function and potentially offer leads for new antivirulence drugs. While classical uncouplers can be used to inhibit proton-powered motors, they also have adverse effects on cells resulting from collapse of the membrane potential. For example the commonly used uncoupler, carbonyl cyanide *m*-chlorophenyl hydrazone (CCCP), is toxic to both prokaryote and eukaryote cells (22). Here, we report that two recently reported 6-substituted amiloride analogues, BB2-50F and HM2-16F(23, 24) (Fig. 1A), inhibit motility in known phenamil-resistant strains and affect both Na^+^- and H^+^-powered flagellar motors.

**Fig. 1:**
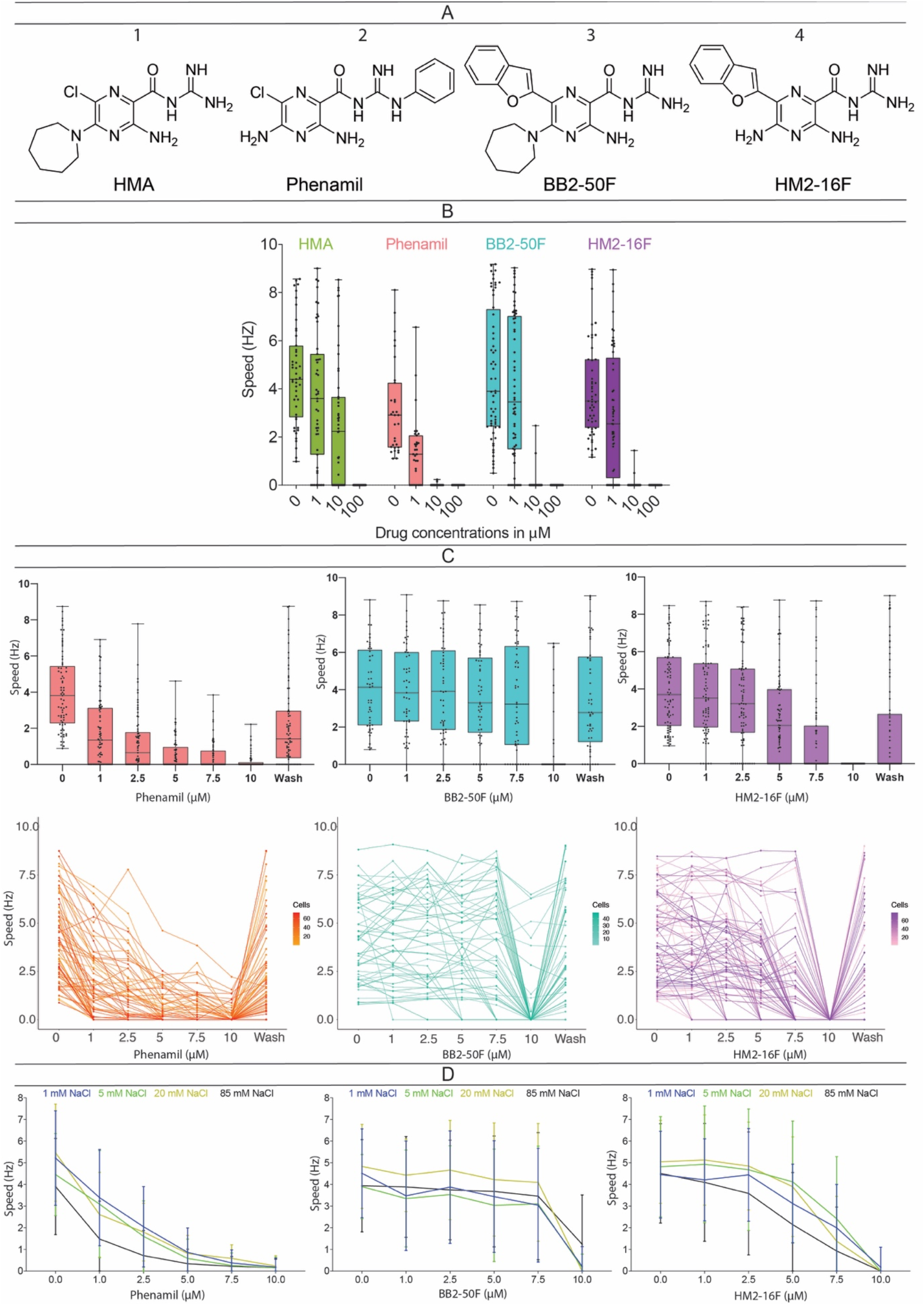
Structures and effects of compounds on bacterial rotation in tethered cell assays. **(A)** Chemical structures of controls 5-*N,N*-(hexamethylene)amiloride (HMA) and phenamil and test compounds BB2-50F and HM2-16F. **(B)** Tethered cell rotation speed (Hz) after treatment with drugs (μM). Cells were *E. coli* SYC35 strain with PomAPotB stators (pSHU1234). Cells were treated with 0, 1, 10 and 100 μM of each compound. Boxes represent the 25^th^ and 75^th^ percentile, with median indicated by dividing line. Whiskers cover the full range from minimum to maximum (n ≥ 30). The lime green bar = HMA, salmon = phenamil, blue = BB2-50F and violet = HM2-16F. **(C)** Dose characterisation of compounds. Cells were treated with 0, 1, 2.5, 5, 7.5 and 10 μM of each compound. The line graphs (bottom) indicate the tracking of individual cells’ change in rotation speed. **(D)** Effects of sodium concentration ([Na^+^]) on inhibitory activity. Cells were treated with 0, 1, 2.5, 5, 7.5 and 10 μM drugs diluted in motility buffer containing different Na^+^ concentrations (1, 5, 20 and 85 mM). Total ionic concentration was held constant and balanced with 85, 80, 65 and 0 mM KCl respectively.

## Results

### Inhibition of motility

We initially evaluated the anti-motility activity of BB2-50F and HM2-16F (Fig. 1A) along with the control compounds 5-(*N*,*N*-hexamethylene)amiloride (HMA) and phenamil using a tethered cell assay (Fig. 1B). We examined the effect of each compound on rotational speed at increasing concentrations (0, 1, 10 and 100 μM) against Na^+^-powered stator, PomAPotB in *E. coli* SYC35 (Δ*motAmotB fliC^sticky^*) strains. Both BB2-50F and HM2-16F inhibited rotation speed in a dose-dependent manner and showed complete inhibition at 10 μM and above (Fig. 1B). The positive control for Na^+^ -inhibition, phenamil showed similar dose-dependency, with complete inhibition at 10 μM (Fig. 1B). HMA was less potent, with 65% of cells still rotating at 10 μM (Fig. 1B), in agreement with previous findings for other 5-substituted amiloride analogues (16).

We further characterised the activity of BB2-50F and HM2-16F alongside phenamil over the concentration range 0-10 μM (Fig. 1C). BB2-50F showed a 2% reduction in the number of rotating cells at 1 μM, 4% at 2.5 μM, 12% at 5 μM, 20% at 7.5 μM and 79% at 10 μM (Fig 1C). HM2-16F showed a 6% reduction in the number of rotating cells at 1 μM, 15% at 2.5 μM, 28% at 5 μM, 62% at 7.5 μM and 100% at 10 μM (Fig. 1C). In comparison, phenamil produced a 27% reduction in the number of rotating cells at 1 μM, 36% at 2.5 μM, 51% at 5 μM, 58% at 7.5 μM and 74% at 10 μM (Fig. 1C). The effect of each compound was reversed following washing with motility buffer.

### Effect on bacterial growth

We measured the growth of *E. coli* SYC35 cells expressing sodium-powered stators (PomAPotB) and proton-powered stators (MotAMotB), and on EHEC and UPEC strains in the presence of 10 μM and 100 μM BB2-50F and HM2-16F (Supplementary Fig. 1). The rate of growth in both strains was not inhibited relative to vehicle controls for either HM2-16F or BB2-50F at 10 μM (Supplementary Fig. 1). However, at 100 μM the compounds differed in toxicity. HM2-16F reduced the growth rate relative to control over the first five hours in both strains, but did not significantly decrease growth at 15 h, while BB2-50F completely inhibited growth at 100 μM.

### Effect of Na^+^ concentration on motility inhibition

We evaluated the inhibition of rotation for BB2-50F, HM2-16F and phenamil while varying the concentration of external sodium to test for Na^+^-competitive effects (17). Compounds were tested at concentrations of 1-10 μM using motility buffers containing 1, 5, 20, and 85 mM Na^+^. No significant differences in inhibition were observed for HM2-16F and BB2-50F at the Na^+^ concentrations tested (Fig. 1D). Inhibition was generally flat for BB2-50F between 0-7.5 μM, with a sharp drop seen at 10 μM, whereas HM2-16F inhibition showed a more gradual effect across the range of concentrations tested. In contrast, an exponential decay was seen for phenamil across the concentration range, with the fitted decay constant indistinguishable at all sodium concentrations (Fig 1D, left).

### Activity against phenamil-resistant variants and the proton powered MotA/MotB complex

To assess whether the inhibition of rotation was affected by point mutations known to confer resistance to phenamil, we tested BB2-50F and HM2-16F against known phenamil-resistant mutant constructs at 1, 10 and 100 μM using the tethered cell assay. Four different stator mutants were tested in SYC35: PomA(D148Y)/PotB, PomA/PotB(P16S), PomA(D148Y)/PotB(P16S) and PomA/PotB(F22Y/L28Q) (68, 69). Phenamil at 10 μM exhibited no significant effect on the resistant strains (Fig. 2A). In contrast, both compounds showed significant inhibition, completely stopping motility at 10 μM in all phenamil-resistant strains (Fig. 2B & C).

**Fig. 2:**
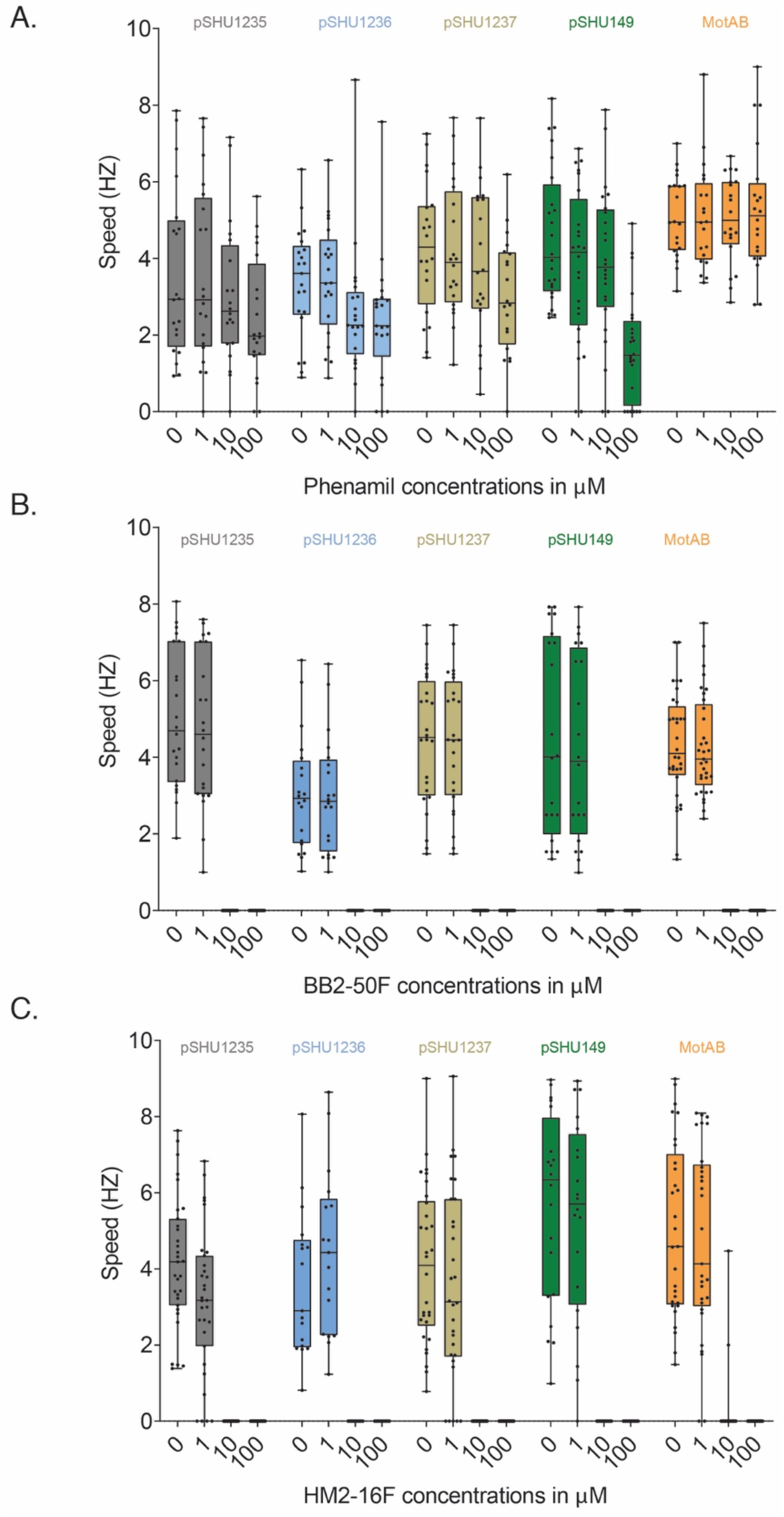
Activity of compounds against phenamil-resistant stators and proton-powered MotAMotB. Tethered cell speed (Hz) in response to 0, 1, 10, and 100 μM of: **A)** phenamil, **(B)** BB2-50F and (**C**) HM2-16F. Cell used for all assays were *E. coli* SYC35 with PomA(D148Y)/PotB (pSHU1235), PomA/PotB(P16S) (pSHU1236), PomA(D148Y)/PotB (P16S) (pSHU1237), PomA/PotB(F22Y/L28Q) (pSHU149) and MotAMotB(pDB-108), respectively. Boxes represent the 25^th^ and 75^th^ percentile, with median indicated by dividing line. Whiskers depict the full range from minimum to maximum (n≥ 20).

To further profile the specificity of the inhibitors, we evaluated BB2-50F and HM2-16F against the proton-powered stator complex (MotAMotB) in SYC35. Both HM2-16F and BB2-50F reduced rotation of the proton-powered motor at 10 μM and completely inhibited rotation at 100 μM (Fig. 2). In contrast, phenamil did not inhibit MotAMotB stators at concentrations up to 100 μM (Fig. 2).

### Inhibition of motility in other bacterial strains

Inhibition of motility by BB2-50F and HM2-16F was tested in three additional bacterial strains using a free swimming assay that measures the speed of individual cells by videography(25). We first demonstrated inhibition of swimming in MotAMotB and PomAPotB expressing cells (Fig. 3AB), followed by three additional strains: uropathogenic *E. coli* (UPEC)(26) (Fig. 3C), an attenuated enterohemorrhagic *E. coli* (EHEC)(27, 28) (Fig. 3D) and a native sodium swimmer *Vibrio alginolyticus* (29) (Fig. 3E). Both BB2-50F and HM2-16F significantly reduced swimming speed at 10 μM in all strains (p < 0.05 in all cases).

**Fig. 3:**
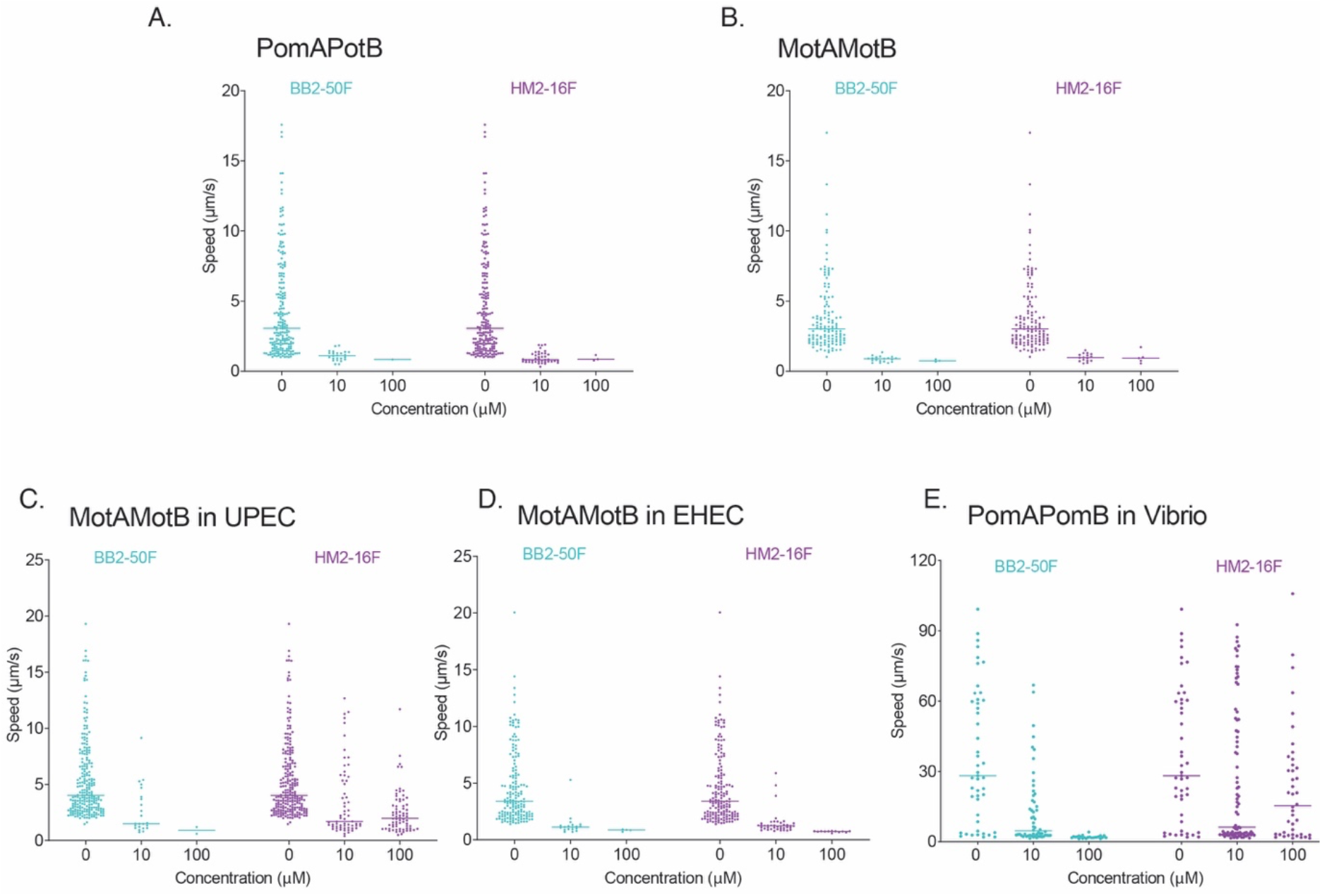
Inhibition of swimming speed by BB2-50F and HM2-16F. Bacterial swimming speed was measured by tracking individual cells using videography in the presence of different concentrations (0, 10, 100 μM) of each compound. Swimming speed of: **(A)** PomAPotB expressed via plasmid in Δ*motAB* strain, **(B)** MotAMotB expressed via plasmid in Δ*motAB* strain, **(C)** MotAMotB in uropathogenic *Escherichia coli* (UPEC) and **(D)** MotAMotB in enterohemorrhagic *Escherichia coli* (EHEC), and **(E)** PomAPomB in *Vibrio alginolyticus*, all following wash with drug. Speeds of individual cells shown in μm/s with the scattered plots, median speed indicated by line (n ≥ 30).

### Characterising mechanism of motility inhibition

The activity of BB2-50F and HM2-16F was measured in a bead rotation assay using 1 μm polystyrene bead attached to sticky flagellar filament of JHC36 (Δ*motAmotB* Δ*cheY fliC^sticky^*) (Fig. 4, Supplementary Fig. 2). This approach reduced the potential for confounding factors that may have affected the tethered cell assay, such as cell-surface adhesion, while also allowing examination of rotation at lower load and at higher speed. Phenamil reduced PomAPotB-driven rotation from 98 ± 7 Hz at 0 μM to 7± 3 Hz at 10 μM (Fig. 4A) but had no effect on the proton-powered MotAMotB (Fig. 4D). In contrast, BB2-50F and HM2-16F reduced the rotation speed of both sodium-powered (PomAPotB) and proton-powered (MotAMotB) motors to zero at 10 μM (Fig. 4BCF).

**Fig. 4:**
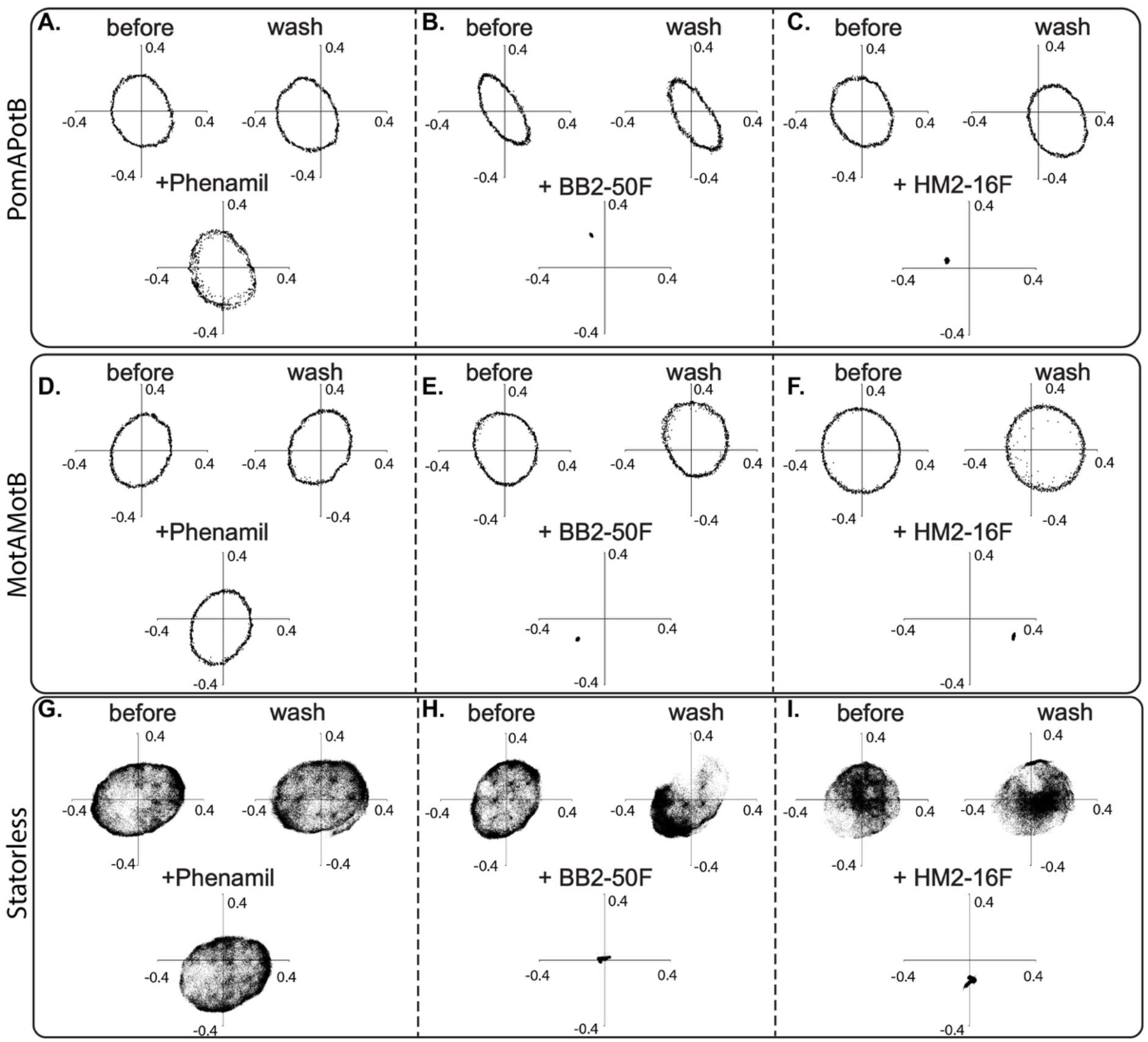
Bead assay results for phenamil, BB2-50F and HM2-16F. **(top)** 1 μm Polystyrene bead assay in *E. coli* strain JHC36 with plasmid expressing PomAPotB stator (pSHU1234) in presence of (A) phenamil, (B) BB2-50F and (C) HM2-16F. Each panel shows three X-Y scatter plots (over 1 s) of the position a bead attached to the rotating flagellar filament of a single cell before exposure to drug, after exposure to drug, and after wash to remove drug. Raw speed vs time and XY vs time plots shown in Supplementary Fig. 2. **(middle)** Bead assay in *E. coli* strain JHC36 with plasmid expressing MotAMotB stator in presence of (D) phenamil, (E) BB2-50F and (F) HM2-16F. **(Bottom)** X-Y position (over 60 s) for *E. coli* strain JHC36 with no stators expressed. The Brownian diffusion of the bead for 60-120 s is shown before and after treatment with (G) phenamil, (H) BB2-50F and (I) HM2-16F, respectively. All drugs washed in as 50 μL of 10 μM drug, washed out with 200 μL of 85MTB. All lengths in μm.

To assess whether the effects of HM2-16F or BB2-50F were due to direct targeting of the stator complex, we measured rotational diffusion of the bead in the presence of drug using the statorless strain. This approach allowed interrogation of whether the compounds were targeting another component of the flagellar motor to inhibit rotation such as the LP ring or bushing. Rotational diffusion for statorless cells remained free in the presence of 10 μM phenamil, indicating no effect on non-stator proteins (Fig. 4G). However, the position of the bead was fixed with minimal rotational or spatial diffusion in the presence of 10 μM BB2-50F (Fig.4H) and HM2-16F (Fig. 4I). We confirmed that rotational diffusion of tethered cells was also restricted in the presence of 10 μM of each compound (Supplementary Fig. 3). Overall, these data suggest a difference in the mode of action of HM2-16F and BB2-50F compared to phenamil, as these compounds appeared to lock the rotational movement of a 1 μm polystyrene bead.

To reduce the possibility of measurements being influenced by lingering surface effects from these comparatively large beads, we reduced bead size and composition, and thus mechanical load, by using a 60 nm gold-nanoparticle attached directly to a straightened hook (30). Given the smaller size of the gold bead, and the straightened hook, this system removed possible bead-cell interactions that may have influenced the earlier polystyrene bead assays. Furthermore, the gold nanoparticle assay allows precise measurement of rotation speed at a load close to zero, providing more direct measure of rotor/stator dynamics free from signal damping caused by viscous drag and mechanical loading (31, 32). For this experiment, we used plasmids expressing proton-powered stator MotAMotB (either pSYC28 or pSYC409) and the sodium-powered stator PomAPotB (pSHU1234) in SHU174 (Δ*motAmotB* Δ*fliC* Δ*cheY flgE_+GSS+3Cys_ fliK2798*). The gold nanoparticle measurements indicated that there was no effect of the compounds at 10 μM on motor rotation driven by either stator type (Fig 5). We further examined the effect of 10 μM of BB2-50F and HM2-16F at low load in statorless cells and observed no measurable difference in rotational diffusion (Fig. 5FG).

**Fig. 5:**
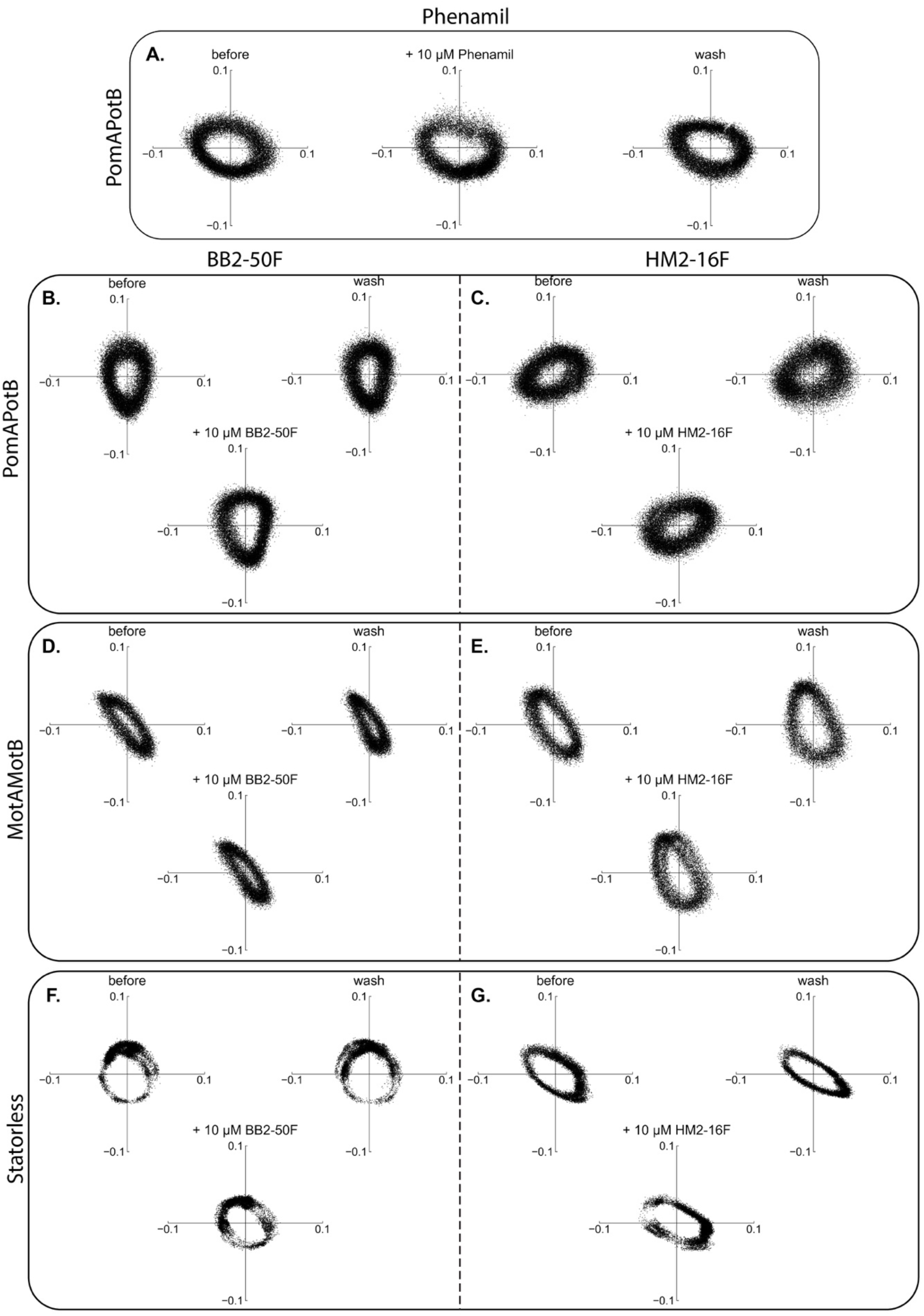
Gold bead nanoparticle assay results phenamil, BB2-50F and HM2-16F. Gold nanoparticle assay was performed in SHU174 *E. coli* strain in presence of 10 μM BB2-50F (left) and HM2-16F (right) for both PomAPotB and MotAMotB stators, with phenamil action on PomAPotB as a positive control (top). (A) Positive control for 10 μM phenamil against rotation driven by PomAPotB. Rotation of full circles is maintained, with slowing of cell speed speed vs time shown in Supplementary Fig. 4). 10 μM of BB2-50F (B) and HM2-16F (C) acting on PomAPotB stators has no effect or gold bead rotation. 10 μM of BB2-50F (D) and HM2-16F (E) acting on MotAMotB has no effect on gold bead rotation. 10 μM of BB2-50F (F) and HM2-16F (G) acting on statorless cells. Rotational diffusion of gold bead tracks circles and is not impeded. Data for PomAPotB, MotAMotB and statorless strainst against 100 μM of compound is shown in Supplementary Fig. 5 & 6. All lengths in μm.

Rotation completely stopped at the 100 μM toxic concentration of BB2-50F for both stator types and for the statorless strains (Supplementary Fig. 4 & 5). In contrast, application of 100 μM of HM2-16F showed mixed effects (Supplementary Fig. 4 & 6). Rotation driven by PomAPotB was not affected, however rotation driven by MotAMotB saw reversible stopping (Supplementary Fig. 6C), irreversible stopping (Supplementary Fig. 6D) and no effect (Supplementary Fig. 6E). Likewise, in statorless strains rotational diffusion of gold beads was both reversibly (Supplementary Fig. 6F) and irreversibly (Supplementary Fig. 6G) stopped.

These results suggest that at non-toxic concentrations of compound (10 μM) both BB2-50F and HM2-16F do not directly target the bushing or the rod of the BFM. Upon observing this, tethered cell assays were repeated in the presence of 0.002% w/v Tween 20 to test whether detergent could remove possible hydrophobic interactions between the bead and cell surface that might underlie the broad anti-motility effect, however no reduction in drug effect was observed (Supplementary Fig. 7). This showed that the effect of the compounds was not removed by a well-validated detergent (33, 34), screening for non-specific interactions between the bead and cell-surface in the presence of each inhibitor. In contrast, phenamil significantly reduced the rotation speed of PomAPotB containing cells in the gold bead assay at 10 μM (Fig. 5A), in accordance with its direct, non-competitive mode of inhibition described previously.

### Fluorescence of HM2-16F

We expected HM2-16F in particular to have improved fluorescence properties relative to phenamil due to the additional benzofuran group conjugated to the pyrazine core. We measured excitation and emission spectra (Supplementary Fig. 8A), and imaged cells in epifluorescence to observe staining with 10 μM HM2-16F (Supplementary Fig. 8B). We further tested HM2-16F using lifetime fluorescence imaging as the quantum yield appeared low (unsuitable for single-molecule imaging). We found that the fluorescence lifetime of HM2-16F was 5.4 ± 0.1 ns, enabling clear resolution against cellular autofluorescence (1.5 ± 0.1 ns) (Supplementary Fig. 8C).

## Discussion

Motility is a potential target for new anti-virulence agents (35). Motility inhibitors are expected to exert less selective pressure than current antibiotics (36), and are typically effective at concentrations lower than those needed for the antibacterial effects of conventional antibiotics (37). This approach may allow for more specific pathogen targeting (e.g. sodium only) as well as lower dosage requirements. Amiloride and its analogues are the best known inhibitors of sodium driven bacterial flagellar motors (16, 17, 38, 39). In this study, we evaluated the anti-motility activity of two new amiloride derivatives, BB2-50F and HM2-16F using four sets of experiments: 1) tethered cell rotation; 2) free-swimming, 3) 1 μm bead rotation, and 4) 60 nm gold bead rotation.

Sodium-driven bacterial rotation was stopped in tethered cell and 1 μm bead assays at 10 μM for BB2-50F, HM2-16F and phenamil, indicating that these compounds show similar potency. The new compounds showed better anti-motility activity at lower concentrations than other reported motility inhibitors for sodium powered motor system, e.g. amiloride and arginine (16, 40). Furthermore, the effect of both BB2-50F and HM2-16F (Fig 1D) appeared to be sodium independent, unlike the Na^+^-competitive parent amiloride (17). In addition to its activity at low concentrations, HM2-16F also showed green fluorescence properties relative to phenamil, enabling the use of epifluorescence and lifetime imaging to measure uptake and release of HM2-16F in treated cells (Supplementary Fig. 8). This finding bolsters its potential applications in, for example, infection diagnostics (41).

BB2-50F and HM2-16F did not affect cellular growth at the motility-blocking concentration of 10 μM (Supplementary Fig. 1), however BB2-50F did inhibit growth at 100 μM. Restoration of motility following washing after dosage with 10 μM of BB2-50F and supported that this dose was non-lethal and that motility could be reversibly restored. In contrast, HM2-16F at 100 μM slowed bacterial growth over the first five hours but was no different from untreated cells after 15 hours (Supplementary Fig. 1). In contrast, BB2-50F was notably toxic at 100 μM, completely inhibiting growth, and likely explaining its consistent anti-motility effect at 100 μM on both stators, in multiple strains, and its irreversible stopping of rotation in the gold bead assay. This indicates a toxic effect of BB2-50F at 100 μM, likely unrelated to any effects on motility proteins. However, it is notable that stopping of rotation in tethered cell assays was seen at a concentration 10-fold lower than this, where no adverse effects on growth were observed in culture (Supplementary Fig. 1).

The dose response of phenamil showed an exponential decrease with increased concentration while BB2-50F, in particular, had a sharp effect appearing between 7.5 μM and 10 μM (Fig. 1B). These results imply that phenamil and both BB2-50F and HM2-16F have different modes of action: both new compounds stopped rotation in phenamil-resistant stator mutants in *E. coli*,and in native stators in other strains, both Na^+^-and H^+^-powered. This property represents an improvement over other reported proton-powered motility inhibitors that are effective only in *E. coli* and require high concentrations (up to 600 μM) (42).

The mechanism of action of these compounds and the reason(s) for their anti-motility activity across multiple bacterial strains is not clear. The tethered cell assay is a good approach for measuring variations in the response of individual cells to treatment and for initial compound screening, however it does not provide positional information for the rotor or assess performance at low load (43). Most importantly, it is also has the largest surface area in motion, which potentially increases its susceptibility to surface based affects such as increased stickiness. To examine the mechanism of inhibition, and to rule out potentially confounding cell-surface effects that may have influenced earlier experiments using tethered cells, we used a 1 μm polystyrene bead assay (44) to examine the effects of BB2-50F and HM2-16F on rotation. We observed that phenamil in some instances slowed, but did not stop, rotation, whereas BB2-50F and HM2-16F stopped rotation and fixed the location of the bead (Fig. 4). We further examined the effect on statorless constructs (Fig. 4G-I) and noted that phenamil had no effect on rotational diffusion in the absence of stators, whereas BB2-50F andHM2-16F locked free rotational diffusion of the bead. The likely explanation is that phenamil directly associates with the stator complex, reducing active torque generation when bound, and thus has no effect in the absence of stators. In contrast, the two compounds appear to jam the motor or adhere the bead to the cell surface, creating transient pauses, or complete stops at higher concentrations.

We used the highest resolution measurement of rotation currently available, a 60-nm gold-nanoparticle affixed directly to a straightened hook (30), to further probe the possible mechanism of action of the compounds. The smaller gold nanoparticle, attached to a straightened hook of ~115 nm length, should not interact with cell surface due to the size of the particle and its distance from the cell membrane. Thus, any observed inhibition would potentially indicate drug binding to the LP ring or bushing of the flagellar motor. However, no inhibition of rotation speed were seen with BB2-50F or HM2-16F at 10 μM in the strains tested using this assay, indicating that the effect of both compounds was likely not due to an on-target interaction with the motor. At high loads there are many stators around a motor, but at low load there should only be a few(31). Our additional measurements of statorless motors at ultra-low load (Fig. 5F & G) saw no change in rotational diffusion in the presence of BB2-50F or HM2-16F at 10 μM. This excludes the possibility that our compounds alter, for example, cell membrane fluidity to interfere with rotation in a stator-dependent manner.

At higher concentrations of 100 μM, BB2-50F irreversibly inhibited motility (Supplementary Fig. 4 & 5). At 100 μM, HM2-16F exhibited the full range of effects: from no effect to reversible and irreversible stopping of rotation (Supplementary Fig. 6) This may indicate that 100 μM is a threshold concentration at which the adhesion effect can influence the gold bead assay. The use of detergent did not change the potency of BB2-50F and HM2-16F in inhibiting rotation (Supplementary Fig. 7).

## Conclusions

We have identified two new amiloride analogues with anti-motility effects at 10 μM concentration in multiple bacterial strains. Whilst the exact mechanism of action of each proved elusive, both compounds were able to inhibit motility in known phenamil-resistant mutants. We have used a wide range of biophysical techniques to rule out possible mechanisms of action: these compounds do not target the stators, the LP ring, or the bushing. Future work can integrate fluorescence measurements with these compounds, and other new derivatives, for targeted drug discovery that aids flagellar research and the development of novel antimicrobials.

## Material and Methods

### Chemical

5-(*N,N*-hexamethylene)amiloride (HMA) (UOW), HM2-16F (UOW) and BB2-50F (UOW) were synthesised as described previously (23, 24). Phenamil methanesulfonate salt (Sigma-Aldrich), Chloramphenicol (Sigma-Aldrich), Dimethyl sulfoxide (DMSO) (Sigma-Aldrich) and D-Arabinose (Sigma-Aldrich).

#### Bacterial Strains and Plasmids and Growth Conditions

The bacterial strains and plasmids used in this study are shown in full in Supplementary Table 1. The primary *E. coli* strains used in this work are SYC35 and RP6894. Pathogenic strains include *Vibrio alginolyticus*, Enterohemorrhagic *Escherichia coli* (EHEC) and Uropathogenic *Escherichia coli* (UPEC). All the strains were cultured in LB broth and LB agar [1% (w/v) Bacto tryptone, 0.5% (w/v) Bacto yeast extract, 0.5% (w/v) NaCl, and 2% (w/v) Bacto agar for solid media] at 37°C. According to the selective antibiotic resistance pattern of the plasmids, chloramphenicol (CAM), ampicillin (AMP) or kanamycin (KAN) were added to a final concentration of 25 μg/mL, 50 μg/mL, and 25 μg/mL, respectively. The *V. alginolyticus* strain, NMB136, was grown in VPG medium (1% (w/v) polypeptone, 0.4% (w/v) K_2_HPO_4_, 3% (w/v) NaCl, 0.5% (w/v) glycerol) and was washed by TMN85 (85 mM NaCl, 215 mM KCl, 50 mM Tris-HCl (pH 7.5), 5 mM MgCl_2_, 5 mM glucose) with appropriate inhibitors.

### Tethered cell assay

We measured the rotation speed of the test strains in the presence of different concentration of test compounds and control drugs using tethered cell assays following a previous protocol with some modifications (45). *E. coli* strain SYC35 (Δ*motAmotB fliC^sticky^*) containing plasmids encoding sodium powered PomAPotB (pSHU1234), proton powered MotAMotB (pDB108) and phenamil resistant PonAPotB (pSHU1235, pSHU1236, pSHU1237, and pSHU1249) were inoculated into TB broth [1% (w/v) Bacto tryptone, 0.5% (w/v) NaCl] containing 0.02% (w/v) arabinose and 25 μg/ml chloramphenicol and were grown overnight (17 h) with 180 rpm at 30 °C. The overnight cultures were sub-cultured with a 50-fold dilution into fresh TB broth and incubated for 5 h with 180 rpm at 30°C to get more motile cells. At OD_600_ ~0.80, the flagella of the cells were sheared by passing the culture multiple times (~35) through a 26G needle syringe. After shearing the flagella, the cells were washed three times with motility buffer [10 mM potassium-phosphate, 10 mM lactic acid, 100 mM NaCl, and 0.1 mM EDTA, pH 7.0]. Cells were then attached on tunnel slides and washed sequentially with motility buffer that contained different concentration test compounds (0, 1, 2.5, 5, 7.5, 10, and 100 μM). According to the experiment purpose we used different concentrations NaCl in the motility buffer. The rotational speed of the cells was observed using phase-contrast microscopy (Nikon) and was recorded at 20 frames per second (FPS) through the 40X objective with a camera (Chameleon3 CM3, Point Grey Research). Rotational motion of the cells was analysed using Lab view 2019 software (National Instruments).

### Free swimming assay

We measured the swimming speed of the test strains in the presence drugs following a previous protocol (46). Briefly, the *E. coli* strains were inoculated into TB broth (1% (w/v) Bacto tryptone, 0.5% (w/v) NaCl) with additional 0.02% (w/v) arabinose and 25 μg/ml chloramphenicol (where required) and were grown overnight (17 h) with 180 rpm at 30°C. NMB136 cells were inoculated in VPG medium (1% polypeptone, 0.4% K_2_HPO_4_, 3% NaCl, 0.5% glycerol) and were grown overnight with 180 rpm at 30°C. For *E. coli* strains, overnight cultures were subcultured with a 50-fold dilution into fresh TB broth and incubated for 5 h with 180 rpm at 30°C. At OD_600_ ~0.80, the cells were washed once with motility buffer (10 mM potassium-phosphate, 10 mM lactic acid, 100 mM NaCl, and 0.1 mM EDTA, pH 7.0). For NMB136 cells, overnight cultures were subcultured with a 100-fold dilution into fresh VPG medium and incubated for 3 h with 180 rpm at 30°C. At OD_600_ ~1.0, the cells were washed twice with TMN85 (85 mM NaCl, 215 mM KCl, 50 mM Tris-HCl (pH 7.5), 5 mM MgCl_2_, 5 mM glucose). Then, cell suspensions were mixed with equivalent volume of the inhibitor solution (at 2x of the intended final concentration) to achieve a final concentration of 10 μM or 100 μM of compound. Finally, 15 μL of cell-inhibitor mixtures were added to the tunnel slide and the swimming speed of the cells was observed using phase-contrast microscopy. For *E. coli* strains: 20 s was recorded at 20 frames per second (fps) through a 20x objective (Nikon) with a camera (Chameleon3 CM3, Point Grey Research). For NMB136 cells: 10 s was recorded at 60 fps through a 20x objective (Olympus) with a camera (DMK21AU618, IMAGING SOURCE). The swimming speed of the cells in both cases was analysed using LabVIEW 2019 software (National Instruments).

#### Growth Curve Assays

Growth assays were performed in the presence of predetermined concentration of test compound to evaluate the effect of the compounds on bacterial growth following the previous protocol (47) with slight modification. *E. coli* strain SYC35 transformed with sodium powered (PomAPotB) and protonpowered (MotAMotB) was used. Briefly, overnight cultures of the test strains with OD_600_ 1.0 were diluted 1:100 in 96-well plates (Corning) with fresh LB broth containing different concentrations of HM2-16F and BB2-50F and incubated at 37°C. Plasmid containing strains (pSHU1234 and pDB-108) were grown in presence of 0.02% (w/v) arabinose and 25 μg/mL chloramphenicol, whereas EHEC and UPEC were grown in LB without arabinose or antibiotic. The OD_600_ were measured in a microplate reader (FLUOstar OPTIMA, BMG LABTECH) every hour for 15 h with a brief shaking interval before each measurement. Measurements in 10 μM were repeated and averaged over three measurements, measurements for 100 μM were repeated and averaged over six measurements.

### Polystyrene bead assay

Briefly, *E. coli* strain JHC36 (Δ*motAmotB* Δ*cheY fliC^sticky^*) transformed with plasmids encoding sodium powered (PomAPotB) and proton powered (MotAMotB) was grown overnight in TB broth and then sub-cultured with a 50-fold dilution into fresh TB broth and incubated for 5 h with 180 rpm at 30°C to get more motile cells. At OD_600_ ~0.80, the flagella of the cells were sheared by passing the culture multiple times (~35) through a 26G needle syringe. After shearing the flagella, the cells were washed three times with motility buffer [10 mM potassium-phosphate, 85 mM NaCl, and 0.1 mM EDTA, 1% DMSO, pH 7.0]. Then, the glass slide was coated with poly-L-lysine (PLL) (Sigma Aldrich) and incubated for 10 mins and washed with motility buffer and flushed with 1 μm polystyrene bead (Polysciences, Inc) solution. The sample was then washed with motility buffer containing test compound (10 and 100 μM) and the rotational speed was recorded and analyzed using Lab view 2019 software (National Instruments).

### Gold-nanoparticle assay

We used the 60 nm gold-nanoparticle assay to get the more accurate flagellar rotation speed at lowered load using the previous protocol (30) with slight modification. Briefly, the SHU174 *E. coli* (Δ*motAmotB* Δ*fliC* Δ*cheY flgE_+GSS+3Cys_ fliK2798*) containing plasmids encoding stators was grown and washed with KPi buffer (10 mM potassium-phosphate, 0.1 mM EDTA, pH 7.0) and prepared as a cell suspension. The cell suspension was later added to the precipitation of gold nano particle (BBI solutions) and incubated for 30 min. After incubation, the cell mixture was washed with KPi buffer and put into PLL coated glass chamber. The sample was then washed with motility buffer (10 mM potassium-phosphate, 10 mM lactate-Na, 67 mM NaCl, 0.1 mM EDTA, 1% DMSO, pH 7.0) containing test compound (10 and 100 μM) and the rotational speed was recorded and analyzed using LabVIEW 2019 software (National Instruments).

### Evaluation of fluorescence property of HM2-16F

The fluorescence of HM2-16F was analysed using fluorescence microscopy. The emission and excitation spectra of the drug was measured using a spectrophotometer (Cary Eclipse). Briefly, the overnight culture of *E. coli* SYC35 strain transformed with pSHU1234 was prepared and washed with motility buffer. Later, the cell suspension was added to PLL-coated glass slide and incubated for 10 mins and again washed with motility buffer. Cells were treated with test compounds by washing with test compound (10 and 100 μM) containing motility buffer.

Finally, drug treated cells were imaged in epifluorescence mode using a Elyra Microscope (Zeiss) using a Plan-Apochromat 63x/1.40 Oil DIC M27 objective, 405 nm illumination and BP495-575 + L750 filter sets. Lifetime imaging was executed on Picoquant MicroTime 200 STED equipped with a PMA Hybrid Series Hybrid Photomultiplier Detector. Illumination was provided by 405 nm laser (Diode 450), with a 60x/1.2 UPlanSApo Water Objective.

## AUTHOR CONTRIBUTIONS

MIM, JB, TI, PR, BJB and MABB designed and executed experiments in, medicinal chemistry, microbiology, microscopy, and rotational measurement. MIM and JL, MJK, BJB, MABB executed initial compound screening. TI and YS designed and executed experiments in rotational measurement with polystyrene beads and gold nanoparticles. MABB and BJB supervised the design, execution and writing of the project. All authors contributed to writing and revision of the manuscript.

## ACKNOWLEDGEMENTS

We would like to thank Tohru Minamino, Yong-Suk Che, Myu Yoshida and Rie Ito for helping a construction of strains and plasmids. We would like to thank Jai Tree for providing the EHEC and UPEC strains and thank Seiji Kojima and Michio Homma for providing the *V. alginolyticus* strain.

## FUNDING

MJK was supported by Australian National Health and Medical Research Council (NHMRC) Project Grant (APP1100432). YS was supported by JSPS KAKENHI (JP18H02475 and JP20K06564), MEXT KAKENHI (JP19H05404) and Takeda Science Foundation. BJB acknowledges salary support from the Illawarra Cancer Carers. MABB was supported by a UNSW Scientia Research Fellowship, a CSIRO Synthetic Biology Future Science Platform 2018 Project Grant, and ARC Discovery Project DP190100497.

## Notes

### Competing Interest Statement

The authors have declared no competing interest.

### Summary of Updates

Rewrote abstract to emphasise correct findings. Altered discussion text. Changed Figure 5 to XY-plot. New Supplementary Figures 2-7.

