## Supplementary Figures 1-8; Supplementary Table 1. for "Novel amiloride derivatives that inhibit bacterial motility across multiple strains and stator types"

**Supplementary Material**

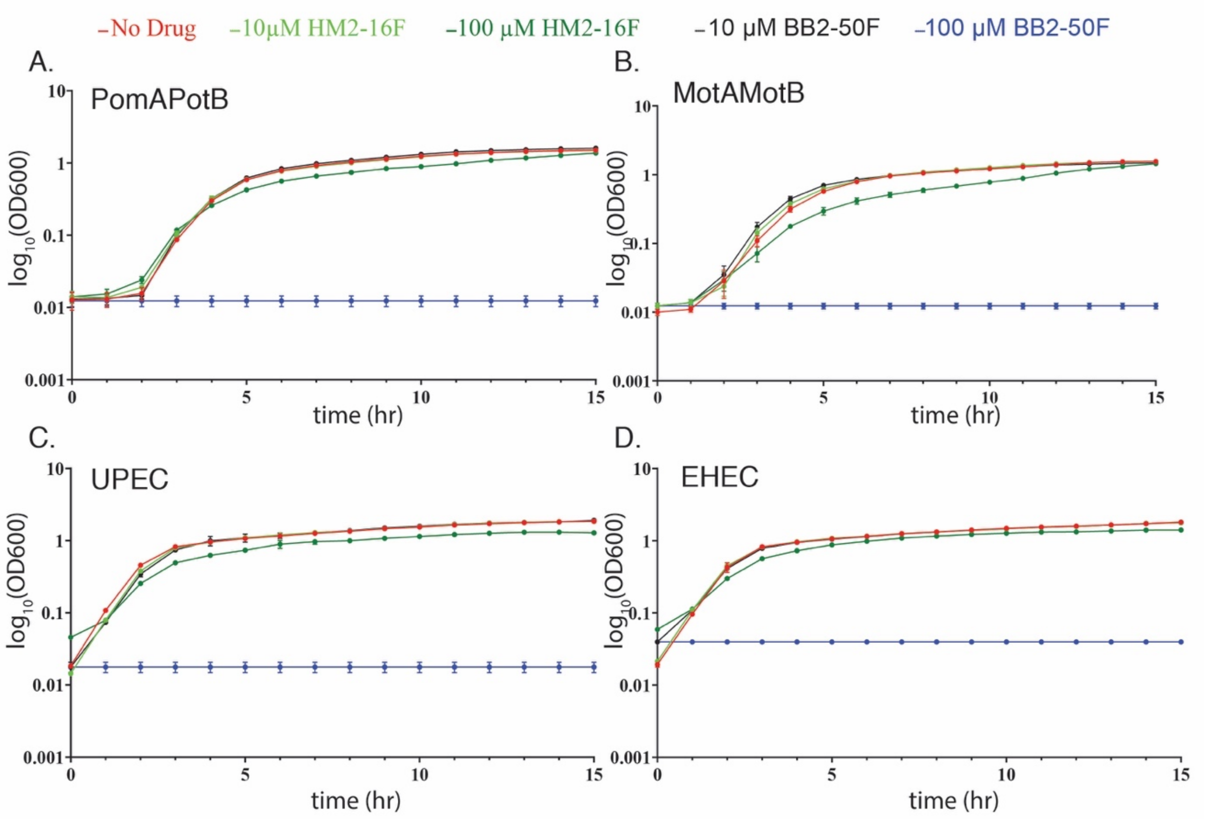

**Supplementary Fig. 1: Effect of compounds on bacterial growth.** Growth curves for *E. coli* strains used in this work in the presence of BB2-50F and HM2-16F using 96 well microtiter plate. (A) SYC35 with sodium-powered (PomAPotB) plasmid, (B) with proton-powered (MotAMotB) plasmid (C) MotAMotB in pathogenic UPEC, (D) MotAMotB in attenuated pathogen EHEC.

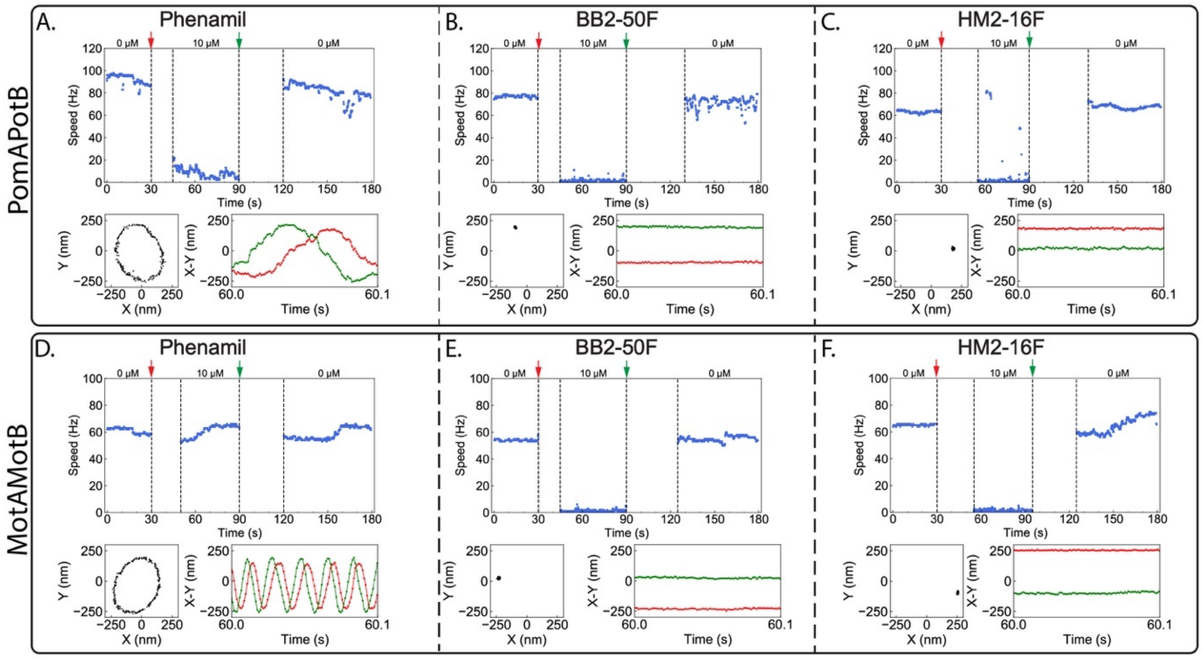

**Supplementary Fig. 2: Bead assay results for phenamil and test compounds. (top)** Bead assay (1 µm polystyrene) in *E. coli* strain JHC36 with plasmid expressing PomAPotB stator (pSHU1234) in presence of (A) phenamil, (B) BB2-50F and (C) HM2-16F. **(bottom)** Bead assay in *E. coli* strain JHC36 with plasmid expressing MotAMotB stator in presence of (D) phenamil, (E) BB2-50F and (F) HM2-16F. Speed versus time over 180 s, with drug washed in at 30 s (red arrow) and out at 90 s (green arrow). Below show X-Y position scatterplot over 0.1 s at 60 s, with X versus time (red) and Y versus time (green) shown adjacently.

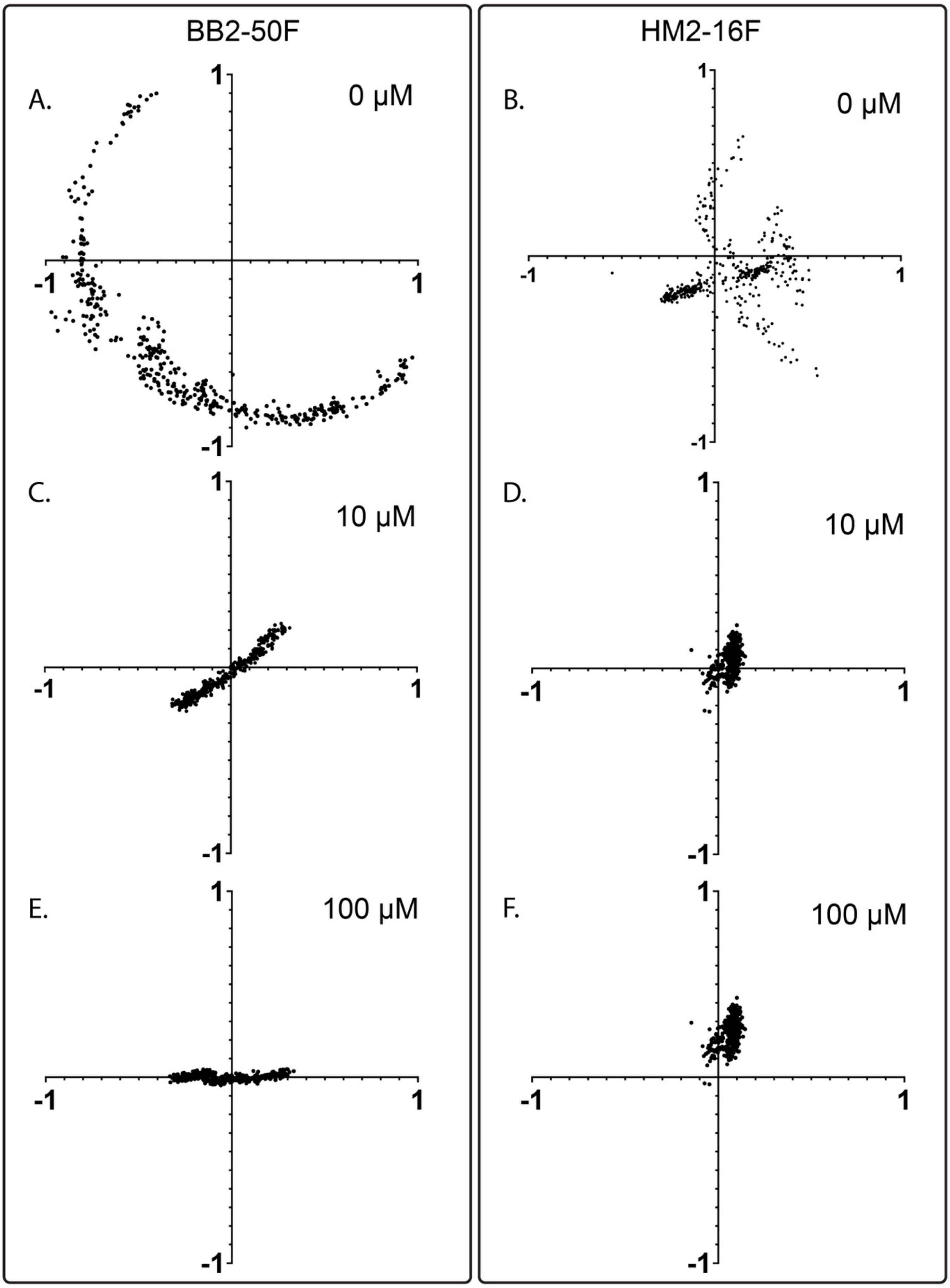

**Supplementary Fig. 3: Effect of compounds on statorless tethered cells.**

Centroid position of sticky filament statorless strain (SYC35) for representative cells in the presence of BB2-50F (left) and HM2-16F (right). There is no stator expression via plasmid. Centroid position for tethered cell diffusing around tether in absence of drug (A/B), in 10 μM of drug (C/D) and in 100 μM of drug (E/F). Cells for BB2-50F and HM2-16F are the same single cell tracked across increasing dosage of drug. Centroid position calculated from 400 frames of video (20 s and 20 fps). All lengths in μm.

26

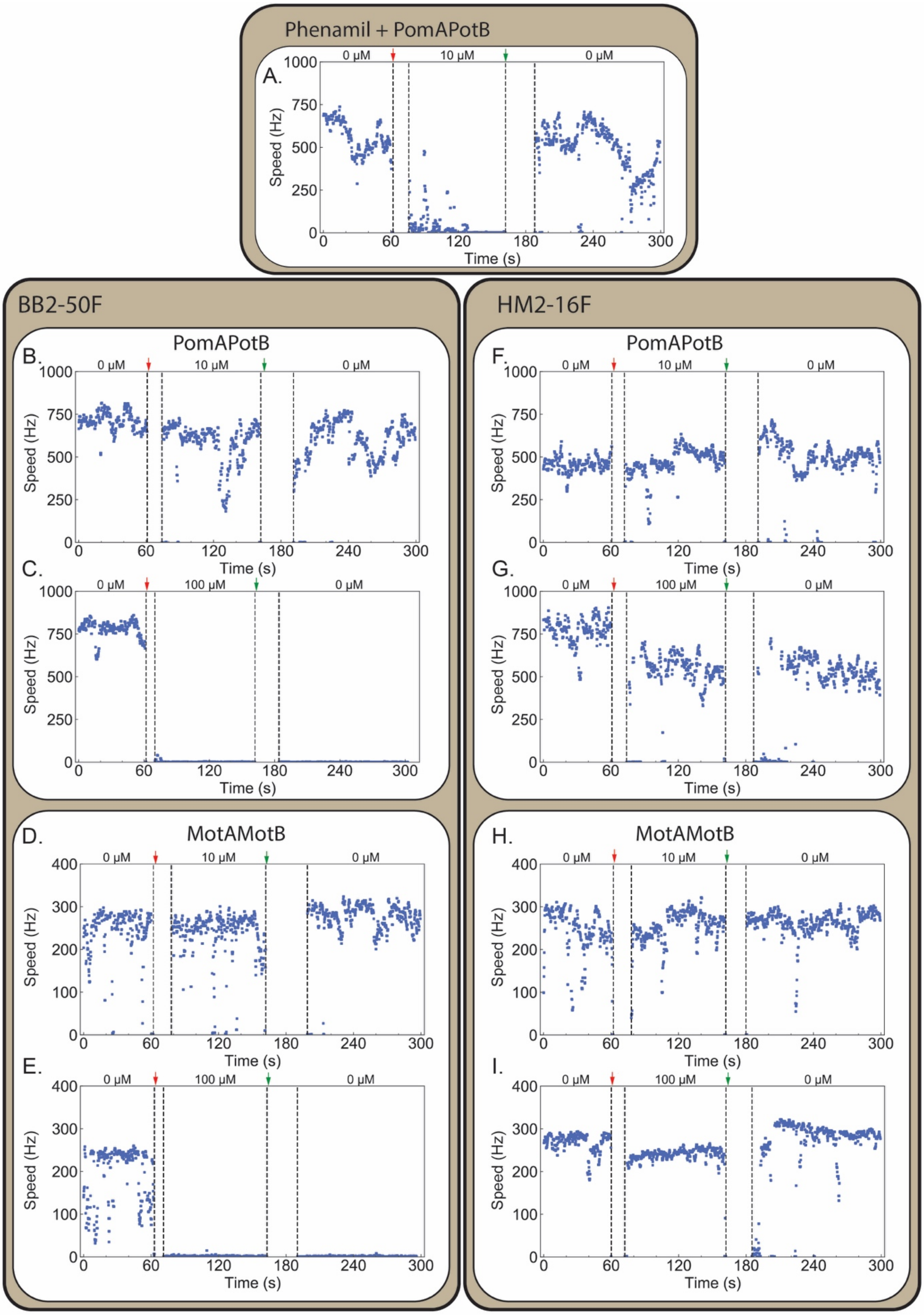
Phenamil + PomAPotB
A.
BB2-50F
HM2-16F
PomAPotB
B.
C.
F.
G.
MotAMotB
D.
E.
H.
I.
0 μM
10 μM
100 μM
Speed (Hz)
Time (s)

**Supplementary Fig. 4: Speed vs time traces for gold bead assay against compounds.** Speed versus time plots for 60 nm gold nanoparticle attached directly to rotating, straightened hook of bacterial flagellar motor before, during and after exposure to drug. Gold nanoparticle assay was performed in SHU174 *E. coli* strain in presence of 10/100 µM BB2-50F (left) and HM2-16F (right) for both PomAPotB and MotAMotB stators, with phenamil action on PomAPotB as a positive control. Red arrow indicates drug flowed in as 50 µL of stated concentration (10 µM or 100 µM), green arrow indicates washout of drug with 200 µL of 67MTB. (A) Positive control for 10 µM phenamil against rotation driven by PomAPotB. Left side: BB2-50F against PomAPotB at (B) 10 µM and (C) 100 µM. BB2-50F against MotAMotB at (D) 10 µM and (E) 100 µM. Right side: HM2-16F against PomAPotB at (F) 10 µM and (G) 100 µM**.** HM2-16F against MotAMotB at (H) 10 µM and (I) 100 µM. Speed vs time traces correspond to X-Y plots shown in Figure 5.

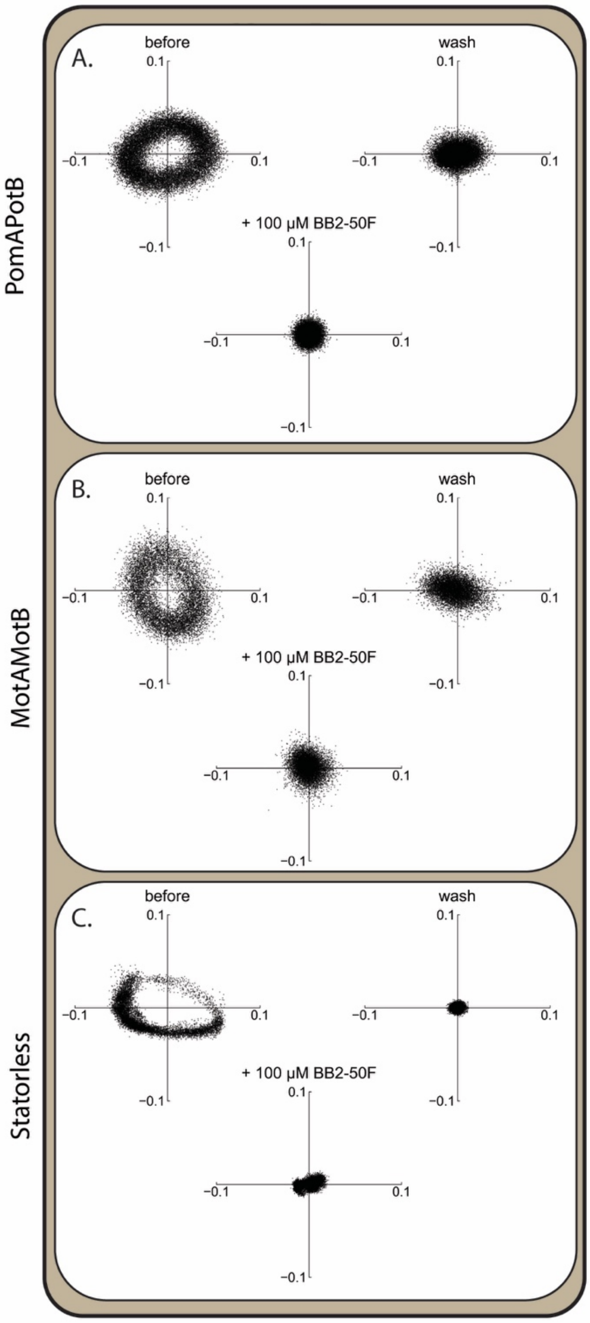

**Supplementary Fig. 5: Effect of 100 µM BB2-50F on rotation of gold bead**. Spatial position in µm of gold bead driven by (A) PomAPotB, (B) MotAMotB and (C) rotational diffusion in statorless cells. In all cases bead motion is irreversibly stopped.

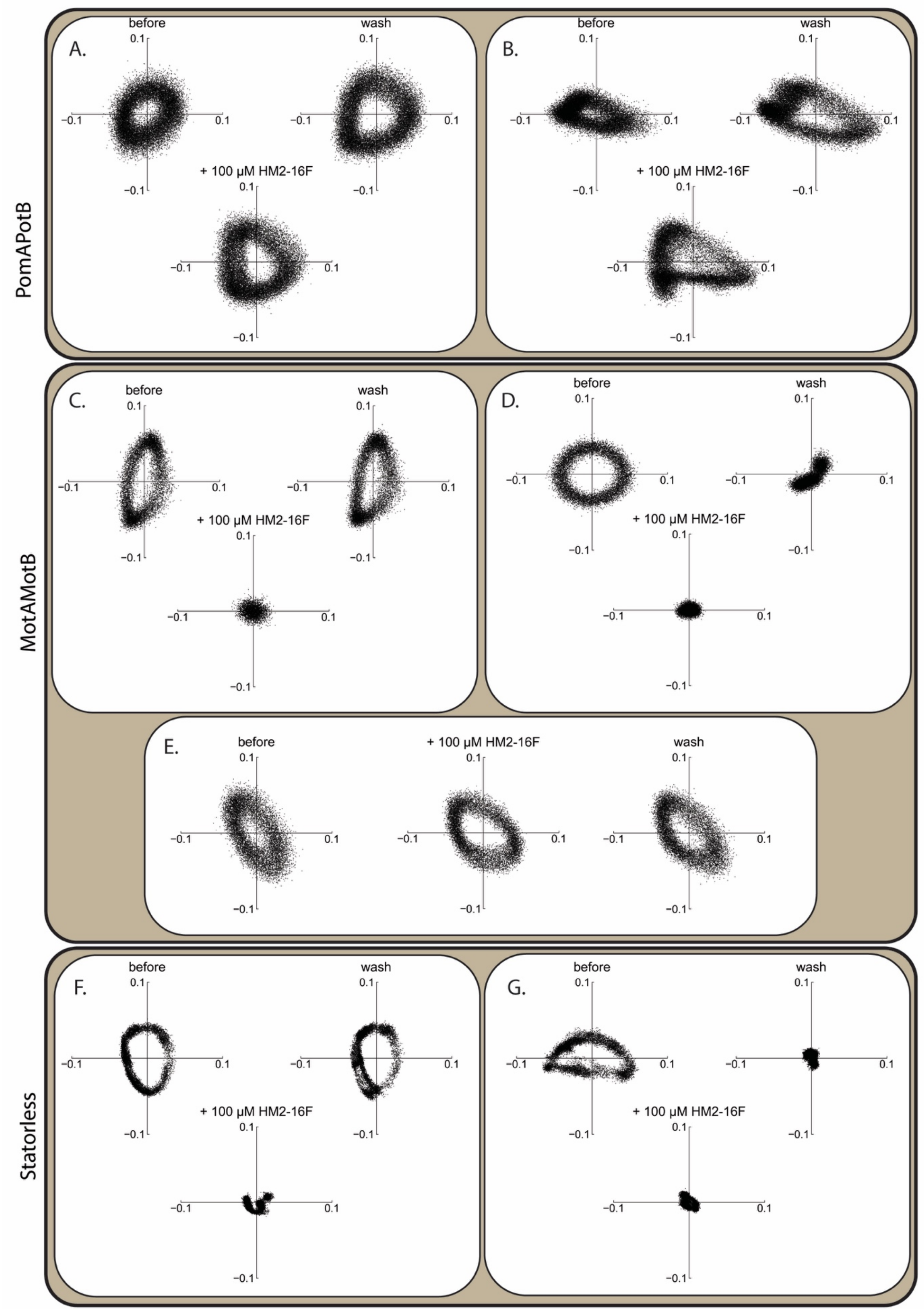

**Supplementary Fig. 6: Effect of 100 µM HM2-16F on rotation of gold bead.** Spatial position in µm of gold bead driven by (A/B) PomAPotB, (C/D/E) MotAMotB and (F/G) rotational diffusion in statorless cells. Effect is varied and representative behaviour in each case is shown. For PomAPotB:

(A/B) no effect; for MotAMotB: (C) reversible stopping after wash, (D) irreversible stopping, (E) no effect; for statorless cells: (F) reversible stopping and (G) irreversible stopping.

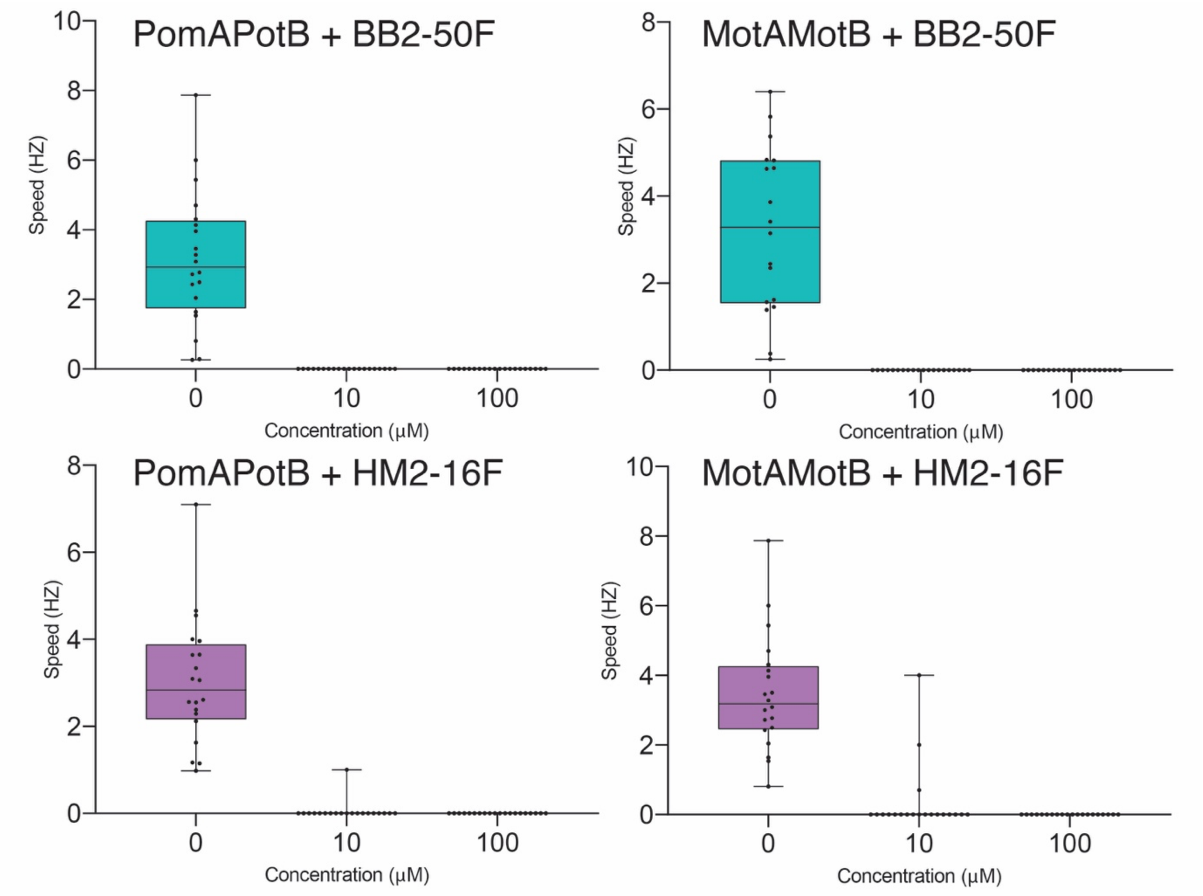

**Supplementary Fig. 7: Drug effect in the presence of Tween-20 on tethered cell assay.** Cell rotation speed (Hz) from tethered cell assay after treatment with 0.002% Tween 20, and BB2-50F and HM2-16F (μM) respectively. The cells were treated with 0, 10 and 100 μM concentration of drug. The cells used were *E. coli* SCY35 strain with pSHU1234 expressing PomAPotB or pDB-108 expressing MotAMotB, respectively (n≥20).

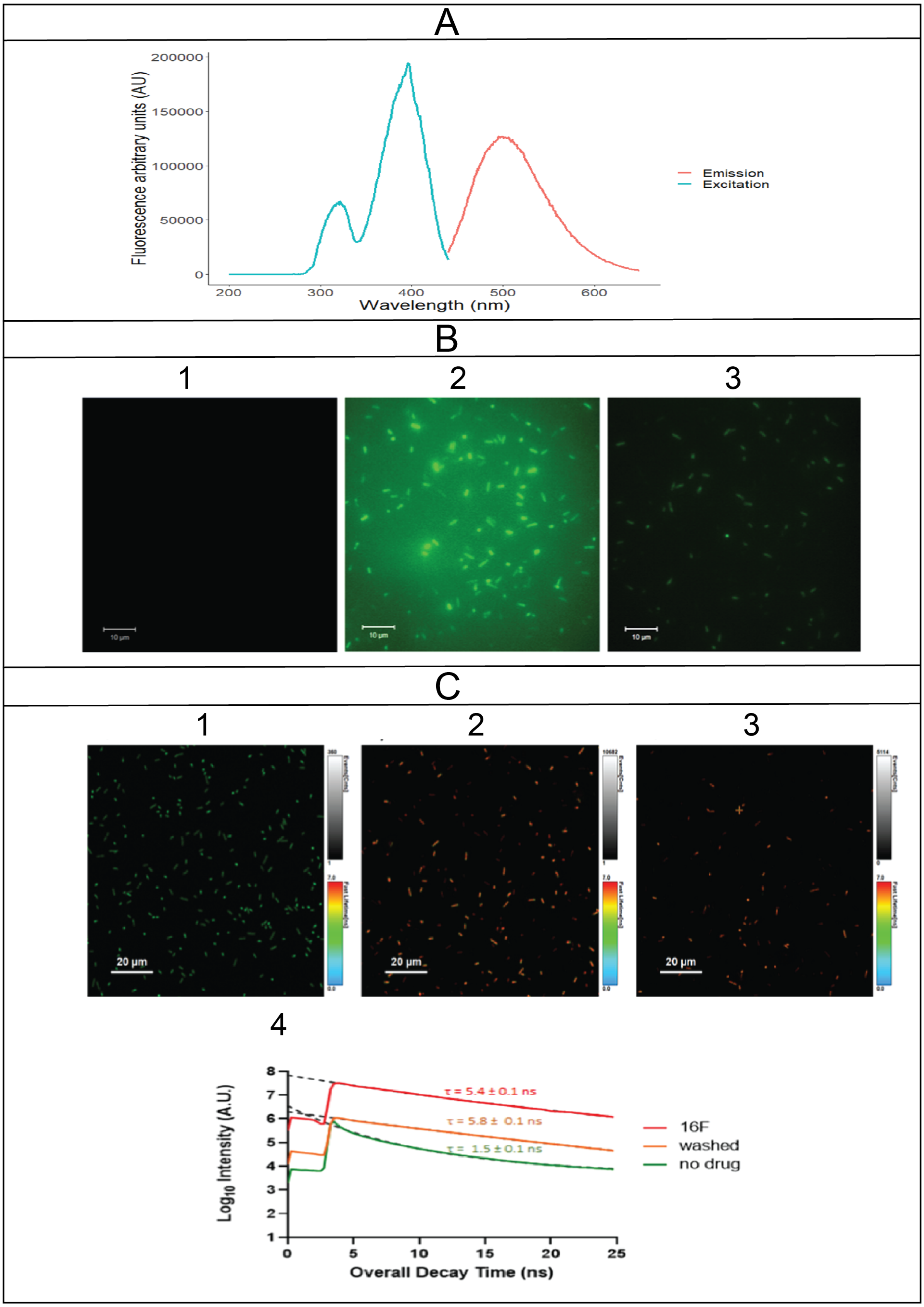
A
Fluorescence arbitrary units (AU)
200000
150000
100000
50000
0
200
300
400
500
600
Wavelength (nm)
Emission
Excitation
B
1
2
3
10 µm
C
1
2
3
20 µm
4
Log10 Intensity (A.U.)
τ = 5.4 ± 0.1 ns
τ = 5.8 ± 0.1 ns
τ = 1.5 ± 0.1 ns
16F
washed
no drug
Overall Decay Time (ns)

**Supplementary Fig. 8: Fluorescence properties of HM2-16F.**

**(A)** Excitation and emission spectra of HM2-16F. Excitation and emission spectra of 10 µM HM2-16F is represented. Cyan represents excitation spectra while orange represents emission spectra. Excitation maximum was 396 nm and emission maximum were 495 nm.

**(B)** Epifluorescence imaging of cells (*E. coli* SYC35 +pSHU1234) treated with HM2-16F. The cells were washed with 10 µM of HM2-16F and then washed with motility buffer. The images taken were an average of 16 images. The minimum pixel intensity was 0 AU and maximum pixel intensity was 192 AU. The images were normalised on ImageJ. (**B-1)** Cells before the treatment. The microscope was unable detect autofluorescence of cells under these parameters. (**B-2)** Cells after 10 µM HM2-16F wash. The mean intensity of cells treated with HM2-16F was 42 ± 4 AU (s.d). (**B-3)** Cells after motility buffer wash. The mean intensity of cells washed with motility buffer was 18 ± 3 AU.

**(C)** Fluorescence lifetime imaging (FLIM) of cell treated with HM2-16F. Fluorescence lifetime imaging of cells (*E. coli* SYC35 + pSHU1234) as it undergoes treatment with HM2-16F. HM2-16F (10 µM) was used to treat the cells and then the drug was washed away using motility buffer. The colour represents the decay rate of fluorescence with red representing a long lifetime and blue representing a short lifetime. (**C-1**) Cells before treatment with the drug. Cells are labelled green. (**C-2**) Cells after treatment with 10 µM HM2-16F. The cells treated with HM2-16F were labelled in orange. (**C-3**) Cells after the drug was washed away with motility buffer. The cells were labelled in red. (**C-4**) The line graph of fluorescence intensity vs overall decay time. The intensity is in $Log_{10}$ values. The green line represents untreated cells, red line represents cells under HM2-16F treatment and orange represents cells washed with motility buffer. The dashed line is the exponential decay curve fitted to each plot. The fluorescence lifetime value ($\tau$) was estimated from the fitting of the exponential decay curve.

**Supplementary Table 1**

| Strain | Description | Reference |
|---|---|---|
| RP6894 | Δ*motAmotB* | J. S. Parkinson (46) |
| SYC35 | Δ*motAmotB fliC*$^{sticky}$ | This work |
| JHC36 | Δ*motAmotB* Δ*cheY fliC*$^{sticky}$ | Inoue et al. 2008 (47) |
| SHU174 | Δ*motAmotB* Δ*fliC* Δ*cheY* *flgE*$_{+GSS+3Cys}$ *fliK2798* | Ishida et al., in preparation |
| **Plasmids** | **Description** | **Reference** |
| pSHU1234 | pBAD33-*pomApotB* | Ishida et al., 2019 |
| pSHU1235 | pBAD33-*pomA*(D148Y)*potB* | This work |
| pSHU1236 | pBAD33-*pomApotB*(P16S) | This work |
| pSHU1237 | pBAD33-*pomA*(D148Y)*potB*(P16S) | This work |
| pSHU149 | pBAD33-*pomApotB* (F22Y/L28Q) | Ishida at. et 2019 |
| pDB108 | MotA and MotB, CAM$^{R}$ | David F Blair |
| pSYC28 | pMMB206-*motAmotB* | Che et al., in preparation |
| pSYC409 | pBAD33-*motAmotB* | Che et al., in preparation |

**Supplementary Movie 1**. Representative swimming video of NMB136 in sequence showing separate slide preparations of: absence of drug, 10 µM BB2-50F, 100 µM BB2-50F, 10 µM HM2-16F, 100 µM HM2-16F. Titles prior to video indicate drug conditions for sample.

**Supplementary Movie 2.** Representative swimming video of SYC36 + pSHU1234 in sequence showing separate slide preparations of: absence of drug, 10 µM BB2-50F, 100 µM BB2-50F, 10 µM HM2-16F, 100 µM HM2-16F. Titles prior to video indicate drug conditions for sample
